# Log D Analysis Using Dynamic Approach

**DOI:** 10.1101/259770

**Authors:** Ganesh Kumar Paul, Prashanth Alluvada, Esayas Alemayehu, M.S. Shahul Hameed, Wasihun Alemayehu, Timothy Kwa, Janarthanan Krishnamoorthy

## Abstract

*Log D* is one of the important parameters used in Lipinski’s rule to assess the druggability of a molecule in pharmaceutical formulations. It represents the logarithm (*log*_10_) of the distribution coefficient (*D*) of a molecule. The distribution coefficient is defined as the ratio of the concentration of the sum of ionized and unionized species of a molecule distributed between a hydrophobic organic phase and an aqueous buffer phase. Since the *pH* affects the ionic state of a molecule, *log D* value (which is dependent on the concentrations of the ionized species) also becomes dependent on *pH.* In this work, the conventional algebraic method is compared with a more generalized ‘dynamic’ approach to model the distribution coefficient of amphoteric, diamino-monoprotic molecule and monoprotic acid in the presence of salt or co-solvent. Recently reported experimental *log D* data of amphoteric molecules such as nalidixic acid, mebendazole, benazepril and telmisartan, were analyzed using both these approaches to show their equivalence.

**GRAPHICAL ABSTRACT:** 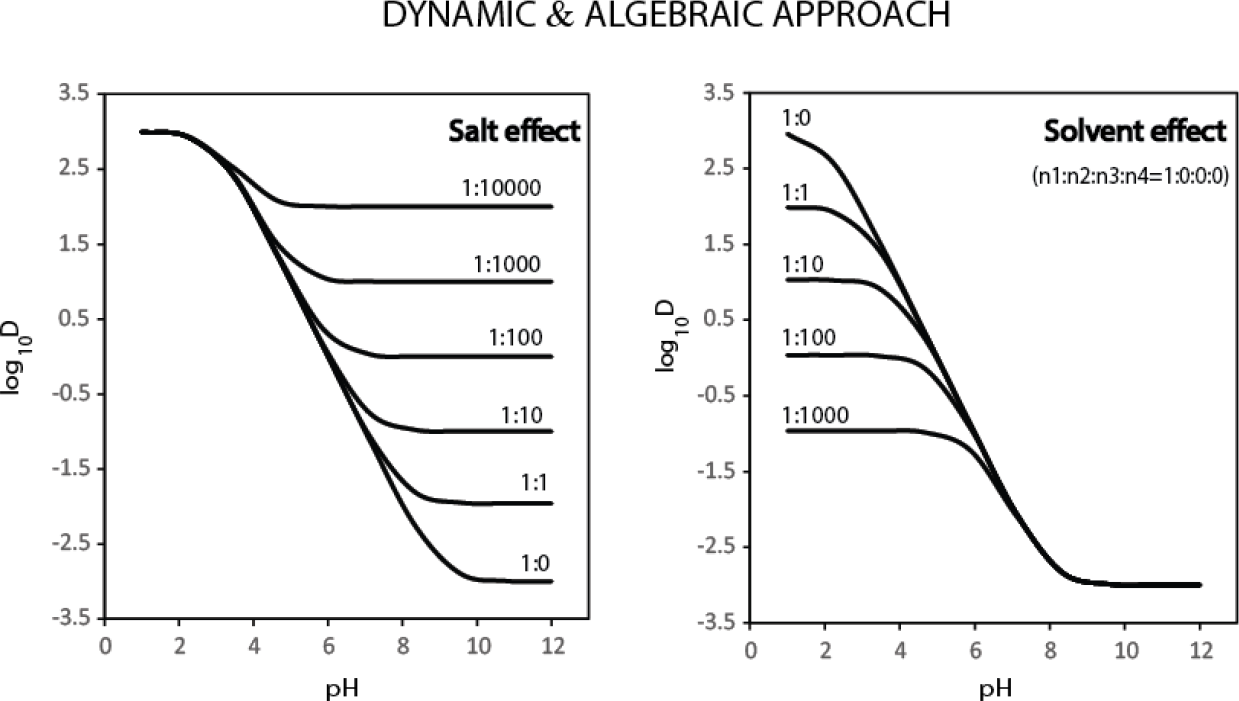

## 1. Introduction

Partition coefficient (*P*) is defined as the ratio of the concentration of a molecule, whether in ionized or unionized form, distributed between a hydrophobic phase and an aqueous phase [1,2,3,4,5]. Consider, a weak monoprotic acid, *HA*, which can exist in two forms such as, unionized (*HA_a_*) and ionized (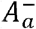) species in an aqueous buffer phase. If the aqueous buffer containing both the species, is equilibrated with a hydrophobic solvent (e.g. octanol), the unionized species in the aqueous phase (*HA_a_*) will get partitioned into the hydrophobic phase (*HA_a_*) with a partition coefficient defined by, *P_HA_* = [*HA*_o_]/[*HA_a_*]. Similarly, the ionized species in the aqueous phase ([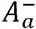]) will get partitioned into the hydrophobic layer ([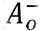]) with a partition coefficient defined as, 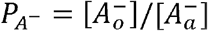. It is less likely for a charged species like *A*^−^, to directly get into an octanol phase, hence, it forms a neutral ion pair with prevalently available cation in the aqueous solution before partitioning into the hydrophobic phase. On the other-hand, the distribution coefficient (*D*), which is dependent on the partition coefficient (*P*), is defined as the ratio of the sum of the concentrations of both ionized and unionized species of a molecule, distributed between the hydrophobic organic phase to the aqueous buffer phase. The distribution coefficient of a weak monoprotic acid, is defined as, 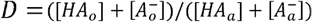. Since the dissociation of the weak monoprotic acid in aqueous phase is dependent on the *pH* value, the distribution coefficient also become dependent on *H*. In an experiment designed to assess the lipophilicity of a molecule, the distribution coefficient (*D*), is measured at different *pH* conditions and the resultant profile of *D*, is fitted to a model, to obtain partition coefficients (*P*), *pK_a_* or *pK_b_* of all the species present in the system [1,2,3,4,5].

The mathematical model to predict the *log D* profile of simple cases such as monoprotic, diprotic, mono-alkaline and amphoteric can be easily derived using algebraic approach[6]. On the other hand, when the effect of salt or solvent has to be taken in to account on the dissociation of monoprotic acid, dynamic approach based modelling could be relatively easier compared to algebraic method [3,5,7,8,9]. In this article, we explicitly, derive the algebraic and dynamic models for amphoteric, di-amino-monoprotic, and monoprotic in the presence of salt or co-solvent [7,8,9]. Further, the *log D* profiles of recently reported amphoteric molecules such as nalidixic acid, mebendazole, benazepril and telmisartan, were analysed to show the equivalence of dynamic approach and algebraic method [10].

## 2. Theory

The algebraic and the dynamic expressions for amphoteric, monoprotic acid in the presence of salt (KCl), co-solvent (DMSO) are explicitly derived in this section. Additional cases such as monoprotic acid (SI.1), diprotic acid (SI.2), monoalkaline (SI.3), diamino-monoprotic amphoteric molecules (SI.7) are detailed in the supplementary information. At the outset, kinetic mechanism that best represent the distribution of the molecule between an aqueous buffer and octanol layer will be outlined. Based on the kinetic mechanism, the algebraic and the dynamic models will be derived.

### 2.1. Amphotheric model for amino acids

#### 2.1.1. Kinetic model for simple amino acids

Consider an amino acid (*NH*_2_ – *R* – *COOH* or *BAH* or *HAB*) containing a weak mono-protic acid (*COOH* or *HA*) and a weak basic/alkaline group (*NH*_2_ or *B*) distributed between an aqueous buffer and an organic hydrophobic solvent (octanol) (Figure 1A) [5,6,11]. In the aqueous phase, the amino acid, [*NH*_2_ – *R* – *COOH*] or [*HAB*], exists in the unionized form [*NH*_2_ – *R* – *COOH_a_*] or [*HAB_a_*], and the ionized forms, 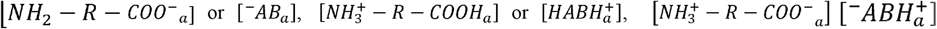. The equilibrium among these four states can be written as (Eqns. 1-5),

**Figure 1.**
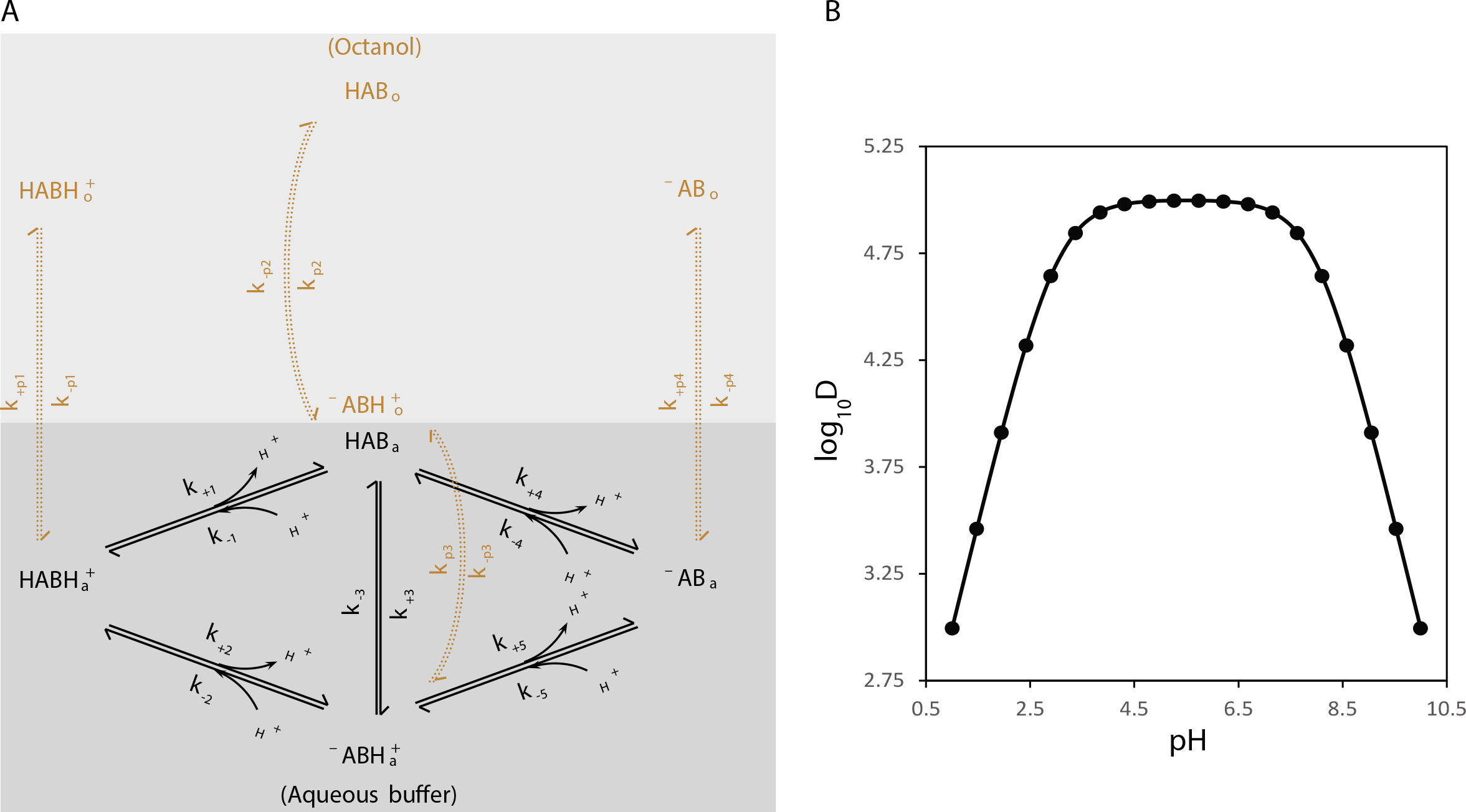
(A) The kinetic model for a simple amphoteric molecule which can exist as four species in aqueous buffer such as, [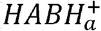], [*HAB_a_*], [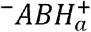], and [^−^*AB_a_*], and gets partitioned into octanol layer as [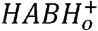], [*HAB_o_*], [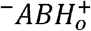], and [^−^*AB_o_*], respectively. (B) shows the simulation of the amphoteric model using algebraic method with equilibrium constants *K*_1_ = *K*_5_ = 10^−8^ and *K*_2_ = *K*_4_ = 10^−3^, partition coefficients, *P*_*HABH*_^+^ = 10^−3^ *P*_−*AB*_ = 10 ^−1^, and *P*_−ABH+_ = *P_HAB_* = 10^5^.

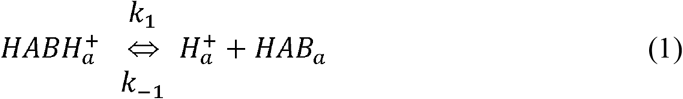

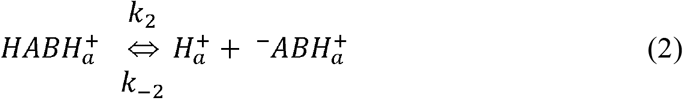

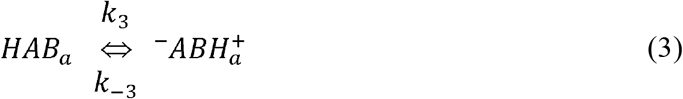

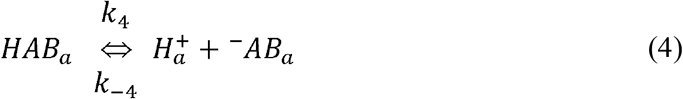

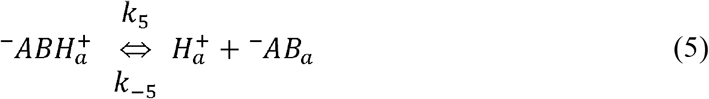

*k*_1_, *k*__1_; *k*_2_,*k*_−2_; *k*_4_, *k*_−4_; *k*_5_, *k*_−5_; are the forward and reverse kinetic rates for the dissociation of the species, such as 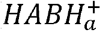 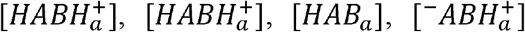, into 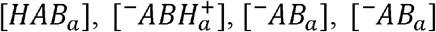, respectively. *k*_3_, *k*_−3_ are the forward and reverse kinetic rates of the tautomerism seen between [*HAB_a_*] and 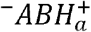, respectively. 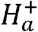, is the proton dissociated from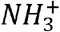 or (*BH*^+^), *COOH* or (*HA*), functional groups. The four species, [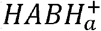], [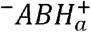], [*HAB_a_*] and [*^−^AB_a_*], can be partitioned into octanol layer as given below (Eqns. 6 - 9),

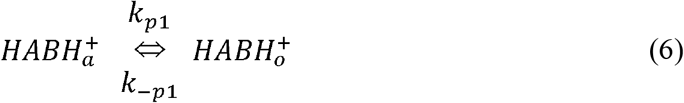

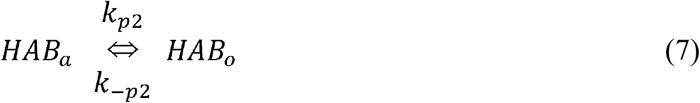

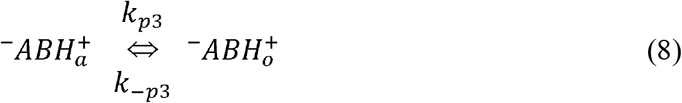

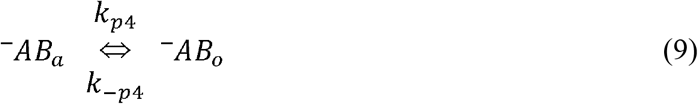

*k*_*p*1_, *k*_−*p*1_; – *k*_*p*2_, *k*−_*p*2_; *k*_*p*3_, *k*−_*p*3_;. *k*_*p*4_, *k*−_*p*4_; are the forward, reverse kinetic rates, respectively, of the partitioning of 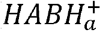, *HAB_a_*, 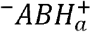, ^−^*AB_a_*, from the aqueous phase into octanol phase, as 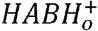, *HAB*_o_, 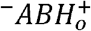, ^−^AB_0_, respectively. The partition of singly charged species such as [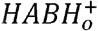], and [^−^*AB_0_*] into the octanol layer can be largely influenced by ion pair formation in the aqueous phase due to the presence of salt. The neutral zwitterions [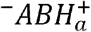], gets easily partitioned into octanol compared to charged species such as [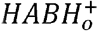] or [^−^*AB_0_*]. Zwitterions exist at a particular *pH* called isoelectric point (*pI*), which is the average of the dissociation constant of its acidic group, *pk_A_*, (*COOH* →; *C00*^−^ + *H*^+^), and the basic group, *pk_B_*, (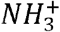 → *NH*_2_ + *H*^+^), i.e. *pI* = (*pk_A_* + *pk_B_*)/2.

#### 2.1.2. Algebraic method for simple amino acids

Based on the above seven equilibriums (Eqns,1,2,4,6-9), seven algebraic equations can be framed in terms of eight species, [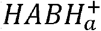], [*HAB_a_*], [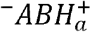], [^−^*AB_a_*], [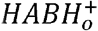], [*HAB_0_*], [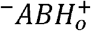], [^−^*AB_o_*]. The distribution coefficient (D) for such a system is defined

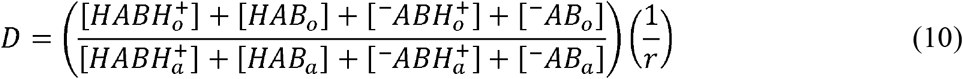

as,
By re-expressing [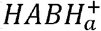], [*HAB_a_*][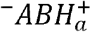], [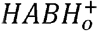], [*HAB*_0_], [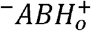], [*^−^AB_0_*], in terms of [^−^*AB_a_*] using the above seven algebraic equations and substituting those expressions into Eqn. 10, we obtain Eqn. 11,

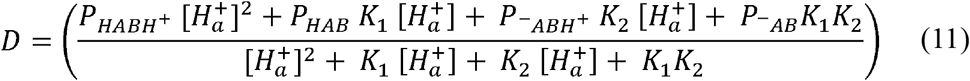

In the above (Eqn. 11), we made a valid assumption on the kinetic mechanism that, *K*_1_ = *K*_5_, and *K*_2_ = *K*_4_. Further, Eqn. 11, will reduce to a diprotic model (SI.4), if *K_1_* = 0, i.e. if the neutral species [*HAB*] is excluded from the kinetic mechanism.

#### 2.1.3. Dynamic method for simple amino acids

Based on the kinetic mechanism (Eqns. 1 - 9), the rate equation (SI.8) can be written for eight species, [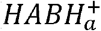], [*HAB_a_*],[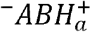], [^−^*AB_a_*], [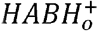], [*HAB*_0_], [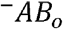] and [^−^*AB*_0_], as follows,

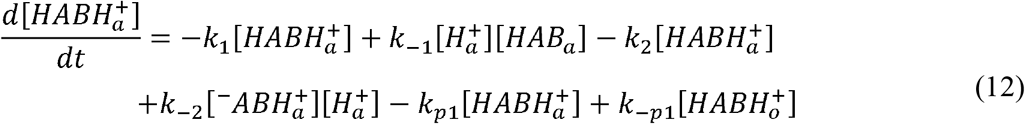

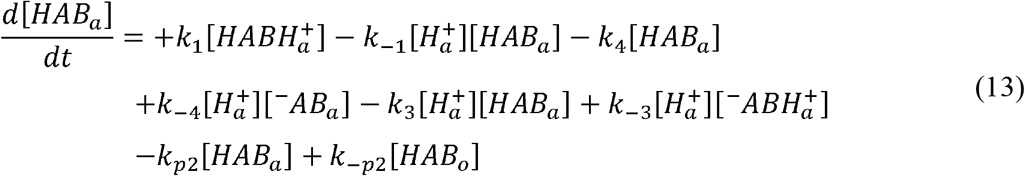

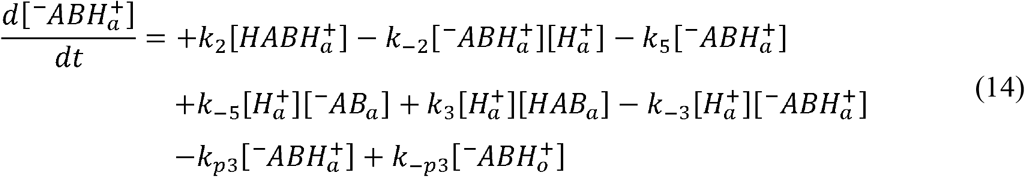

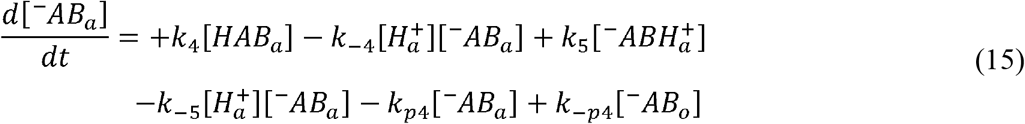

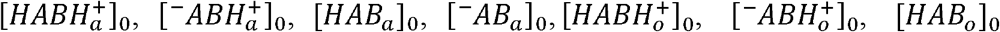

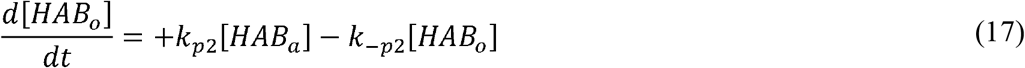

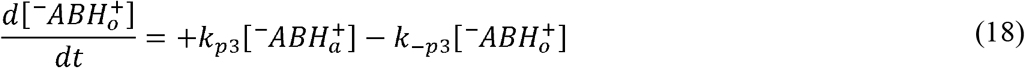

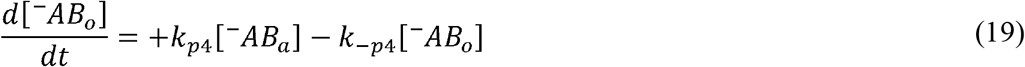

By numerically integrating the above set of coupled differential equations, Eqns 12-19, we obtain the concentration of eight species, [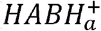], [*HAB_a_*], [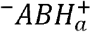], [^−^*AB_a_*], [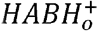], [*HAB_o_*], [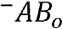], and [^−^*AB_o_*] at different time points. The resultant concentrations at equilibrium, can be substituted into the Eqn. 10, to obtain the distribution coefficient. In the above model, [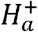] or *pH*, is assumed to be constant with respect to time, due to the use of appropriate buffers in *log*_10_*D* experiments that stabilizes the *pH* of the aqueous phase. To simulate the *log*_10_*D* profile in Figure 1B, the parameters such as *k*_1_ *k*_−1_, *k*_2_, *k*_−2_, *k*_3_, *k*_−3_, *k*_4_, *k*_−4_, *k*_5_, *k*_−5_, *k*_*p*l_, *k*_−*p*l_, *k*_*p*2_, *k*_−*p*2_, *k*_*p*3_, *k*_−*p*3_, *k*_*p*4_, *k*_−*p*4_, were set to 10^−8^, 1.0, 10^−3^, 1.0, 10^−(8+3)/2^, 1.0, 10^−3^,1.0, 10^−8^,1.0, 10^−3^, 1.0, 10^5^, 1.0, 10^5^, 1.0, 10^−1^, 1.0, respectively. The initial concentrations of all the eight species 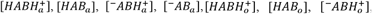 and [^-^*AB*_o_]_0_, were set to 1.0,0,0,0,0,0, 0,0, respectively and the *pH*, was varied linearly between 1 and 10.

### 2.2. Monoprotic acid with salt (KCl)

#### 2.2.1. Kinetic model for monoprotic acid with salt (KCl)

Consider the distribution of a weak mono-protic acid (*HA*) between an aqueous solvent and an organic hydrophobic solvent (octanol) in the presence of a potassium chloride (KCl) (Figure 2A) [3, 5, 12, 13].

In aqueous phase, the weak acid, [*HA*], exists in the unionized form [*HA_a_*] and ionized form [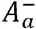]. The equilibrium between these two states can be written as (Eq. 20),

**Figure 2.**
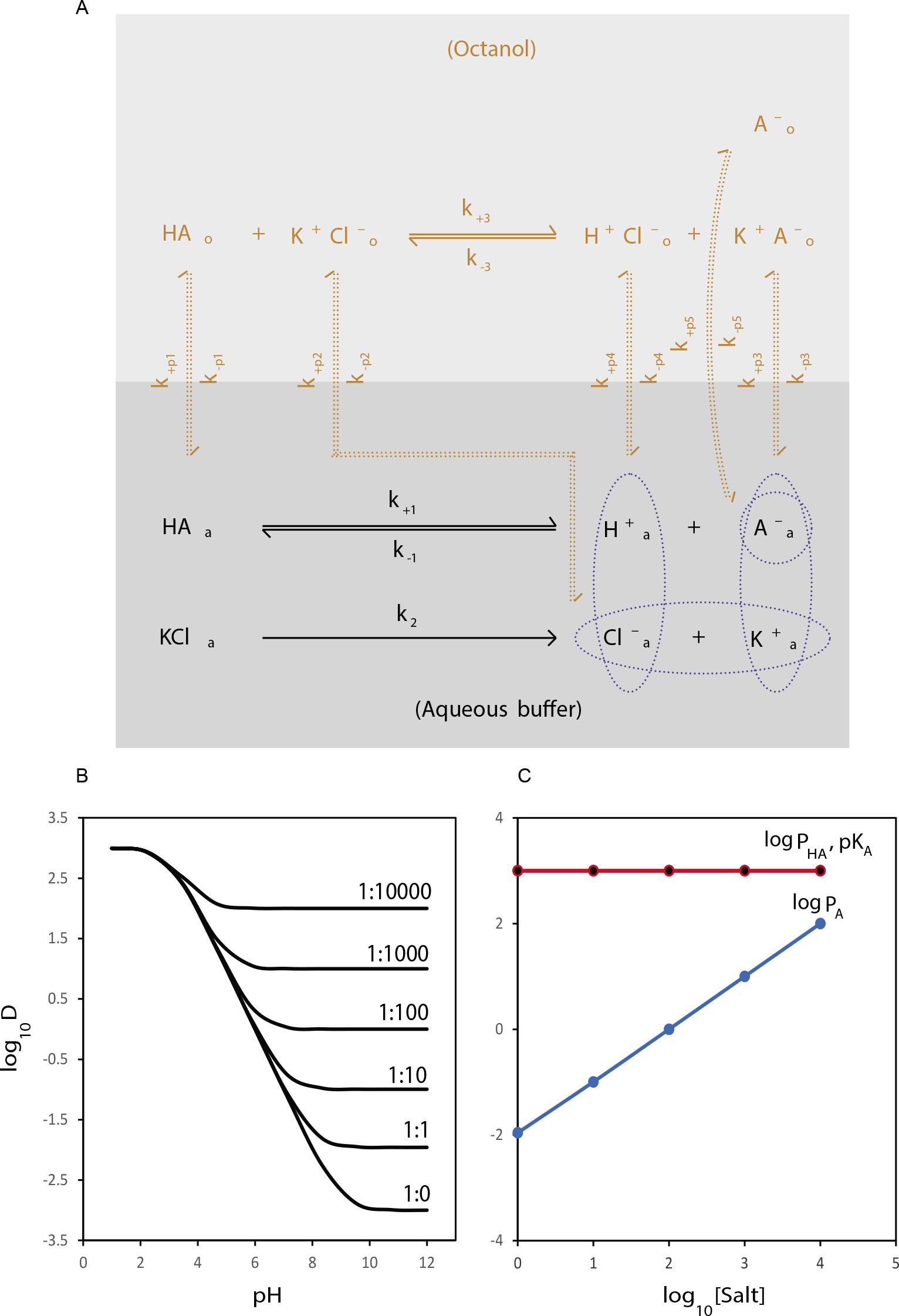
(A) The kinetic model for a monoprotic acid in the presence of a salt. In this mechanism, there exist a reversible equilibrium between *HA_a_*, and 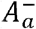 in aqueous phase and a reversible equilibrium between *HA_o_* and 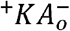 in the octanol phase. The formation of 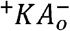 in the octanol layer is mediated through the partitioning of the ion pairs 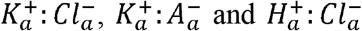 from the aqueous to octanol layer. (B) Shows the *log*_10_*D* profiles of the monoprotic acid when the ratio of monoprotic acid to salt is varied between 1:0 to 1:10000. (C) shows how the *log*_10_ *P_HA_*, *log*_10_, *P_A_*, *pK_A_* of the monoprotic acid varies as the salt concentration increases, from 0 to 10000, plotted here in *log*_10_ scale. It is clear that only *log*_10_ *P_A_* increases linearly with increase in salt concentration, whereas, *log*_10_ *P_HA_* and *pK_A_*, remains constant. (Though not shown, the plot of nonlogarithmic form of *P_A_* vs [*Salt*] is also linear). In the simulation, the *P_HA_*, *P_A_* and *K_A_* of the monoprotic acid were set to 10^3^, 10^-3^ and 10^3^ respectively, the concentration of monoprotic acid was set to 1, and the concentration of salt was set to 0, 1, 10, 100, 1000,10000 to simulate different *log*_10_*D*, profiles. The resultant profiles were fitted to the monoprotic model, 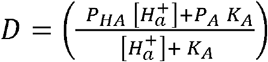, to obtain the apparent 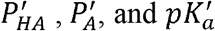, the logarithmic form of which are plotted against the logarithm of salt concentration in (C)

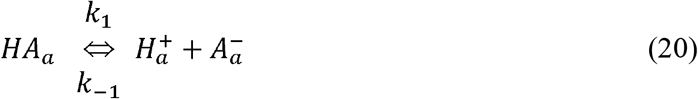

*k*_1_, *k*_−1_ are the forward and reverse kinetic rates for the dissociation of the weak acid, [*HA*], respectively. The salt KCl dissociates completely in aqueous solution to form [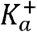] and [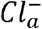] with a forward kinetic rate of *k_2_* (Eqn. 21).

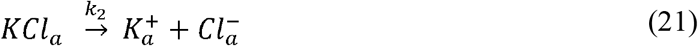

The unionized species, [*HA_a_*] can be partitioned into octanol layer with the forward and reverse kinetic rates as *k*_*p*1_, *k*_−*p*1_, as given below (Eqn. 22),

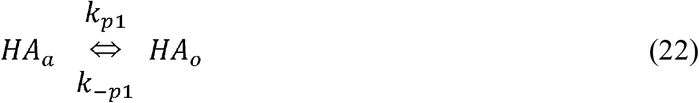

The dissociated [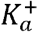] and [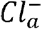] gets partitioned directly into octanol with the forward and reverse kinetic rates as *k*_*p*2_, *k*_−*p*2_, respectively (Eqn. 23).

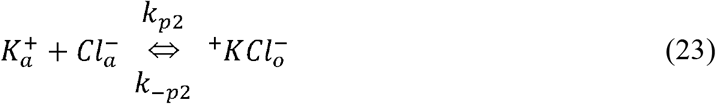

The ionized species [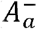], would require its negative charge be neutralized before partitioned into octanol layer through the prevalent cation, 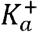, coming from the dissociation of KCl. The forward and reverse kinetic rates of the partition of [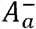]: [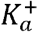] are *k*_*p*3_, *k*_−*p*3_, respectively, as given below (Eqn. 24),

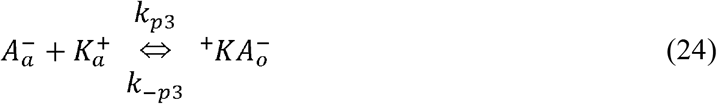

Additionally, as seen with the monoprotic acid (SI.1), it is also possible for a fraction of the ionized species [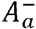], to get directly partitioned into the octanol layer without ion-pair formation as [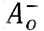], whose forward and reverse kinetic rates are given as *k*_*p*5_, *k*_−*p*5_, respectively, (Eqn. 25),

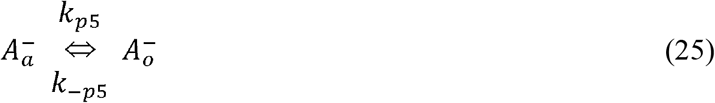

The [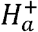] ions (arising from the dissociation of *HA_a_*, (Eq. 20)) and [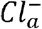] (from [*KCl_a_*]) gets partitioned into octanol with the forward and reverse kinetic rates as *k*_*p*4_, *k*_−*p*4_, respectively (Eqn. 26).

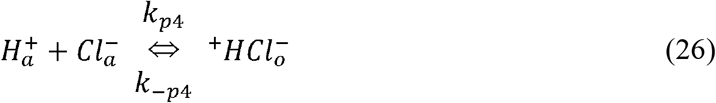

Finally, all the four species *HA_o_*, 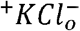, 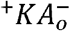, 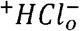, partitioned into the octanol layer undergo a dynamic equilibrium whose forward and reverse kinetic rates are given as *k*_3_, *k*_−3_, respectively (Eqn. 27).

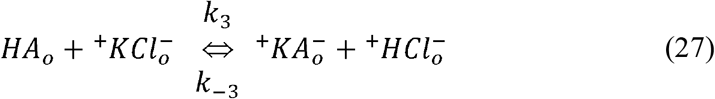

#### 2.2.2. Algebraic method for monoprotic acid with KCl

Applying Le-Chaterlier’s principle, on the above six equilibriums (Eqns. 20, 22, 23, 25, 26 & 27), and considering the mass balance for [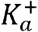], [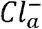], and [*HA_T_*], we obtain the equilibrium concentrations of [*HA_o_*], [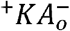], [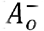], [*HA_a_*] and [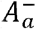] (SI.5). The distribution coefficient (D) of the monoprotic system in the presence of salt is defined as,

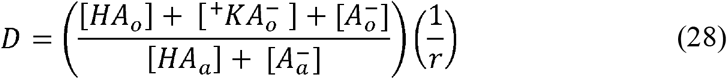

By substituting the analytical expressions of [*HA_o_*], [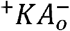], [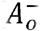], [*HA_a_*] and [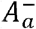] into Eqn. 28, we obtain,

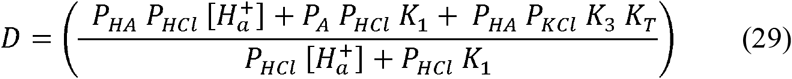

To simulate the *log*_10_*D* profile in Figure 2B, the parameters such as *K*_1_, *K*_3_, *P_HA_*, *P_A_*, *P_Hcl_*, *P_KCI_*, were set to 10^−3^, 10^−8^, 10^3^, 10^−3^, 10^−15^, 10^−15^, respectively. *K_T_* was set to, 0, 1, 10, 100, 1000 and 10000 M, to simulate 6 profiles and the *pH*, was varied linearly, between 1 and 10. If we set either, *K*_3_ or *P_KCL_* = 0, the above Eqn. 29 will reduce to that of a model of simple monoprotic acid.

#### 2.2.3. Dynamic method for monoprotic acid with KCl

Based on the kinetic mechanism (Eqns. 20 - 27), the rate equation can be written for the ten species, [*HA_a_*], [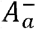], [*KCl_a_*], [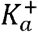], [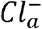] [*HA_o_*], [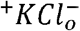], [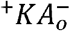], [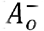], and [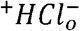], as follows,

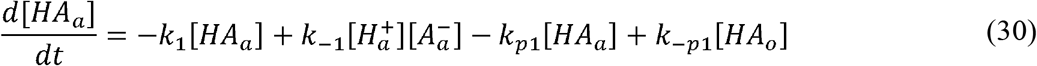

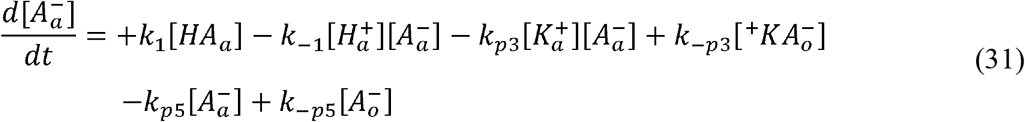

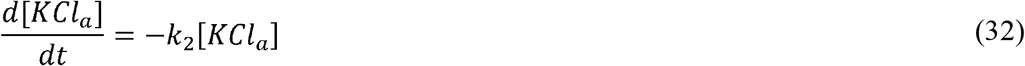

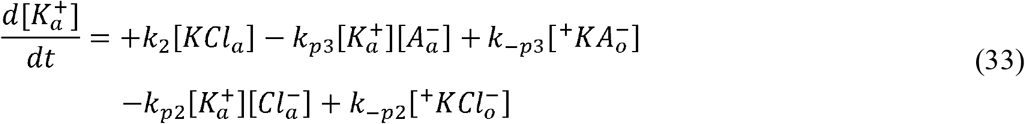

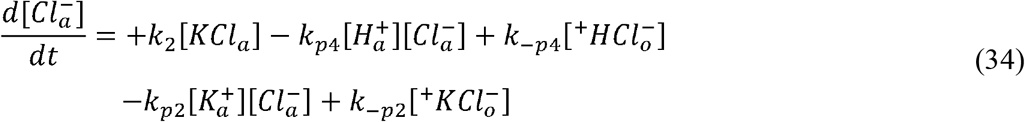

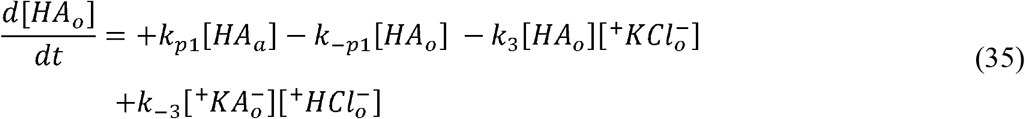

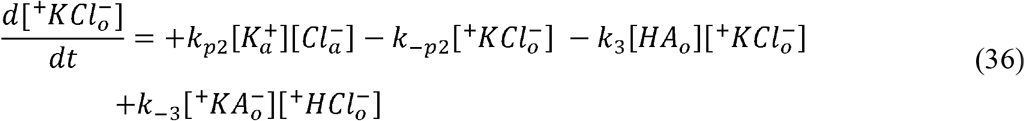

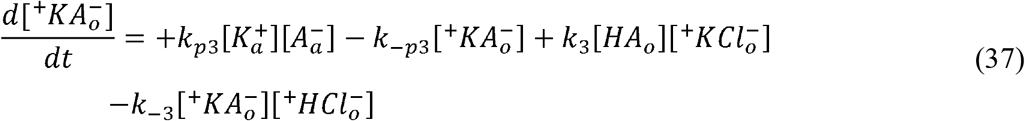

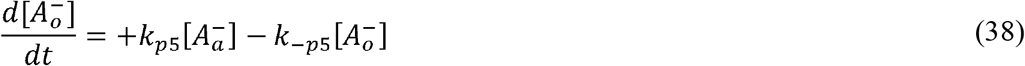

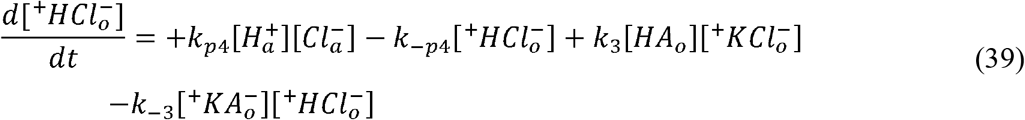

By numerically integrating the above set of coupled differential equations, Eqns 30 - 39, we obtain the concentration of ten species, [*HA_a_*], [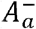], [*KCl_a_*], [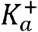], [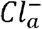], [*HA*_o_], [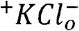], [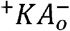], [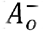] and [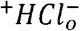], at different time points. The resultant concentrations of [*HA_a_*], [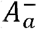], [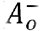], [*HA*_o_], [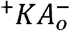] at equilibrium, can be substituted into the Eqn. 28, to obtain the distribution coefficient. In the above model, [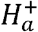] or *pH*, is assumed to be constant with respect to time, because of the usage of appropriate buffers in *log*_10_*D* experiments to stabilize the pH of the aqueous phase. To simulate the *log*_10_*D* profile in Figure 2B, the parameters such as *k*_1_, *k*_−1_, *k*_2_, *k*_3_, *k*_−3_, *k*_*p*l_, *k*_−*p*l_, *k*_*p*2_, *k*_−*p*2_, *k*_*p*3_, *k*_−*p*3_, *k*_*p*4_, *k*_−*p*4_, *k*_*p*5_, *k*_−*p*5_ were set to 10^−3^, 1.0, 10^15^ (any large value, >10^2^), 10^−8^, 1.0, 10^3^, 1.0, 10^−15^, 1.0, 10^−2^, 1.0^−15^, 1.0, 10^−3^, 1.0 respectively. The initial concentrations of the following nine species, 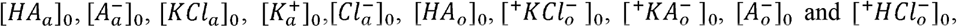 were initialized to 1.0, 0, *K_T_,0,0,0*, 0,0,0,0 M respectively. Six different simulations were performed by varying the initial concentration of *K_T_*, as, 0, 1, 10, 100, 1000, 10000 M, and the *pH* was varied linearly between 1 and 12.

### 2.3. Monoprotic acid in the presence of co-solvent

#### 2.1.3. Kinetic model for monoprotic acid in the presence of co-solvent

Consider the distribution of a weak mono-protic acid (*HA*) between an aqueous solvent and an organic hydrophobic solvent (octanol) in the presence of co-solvent (*S*) such as DMSO (Figure 3A) [3, 4].

**Figure 3.**
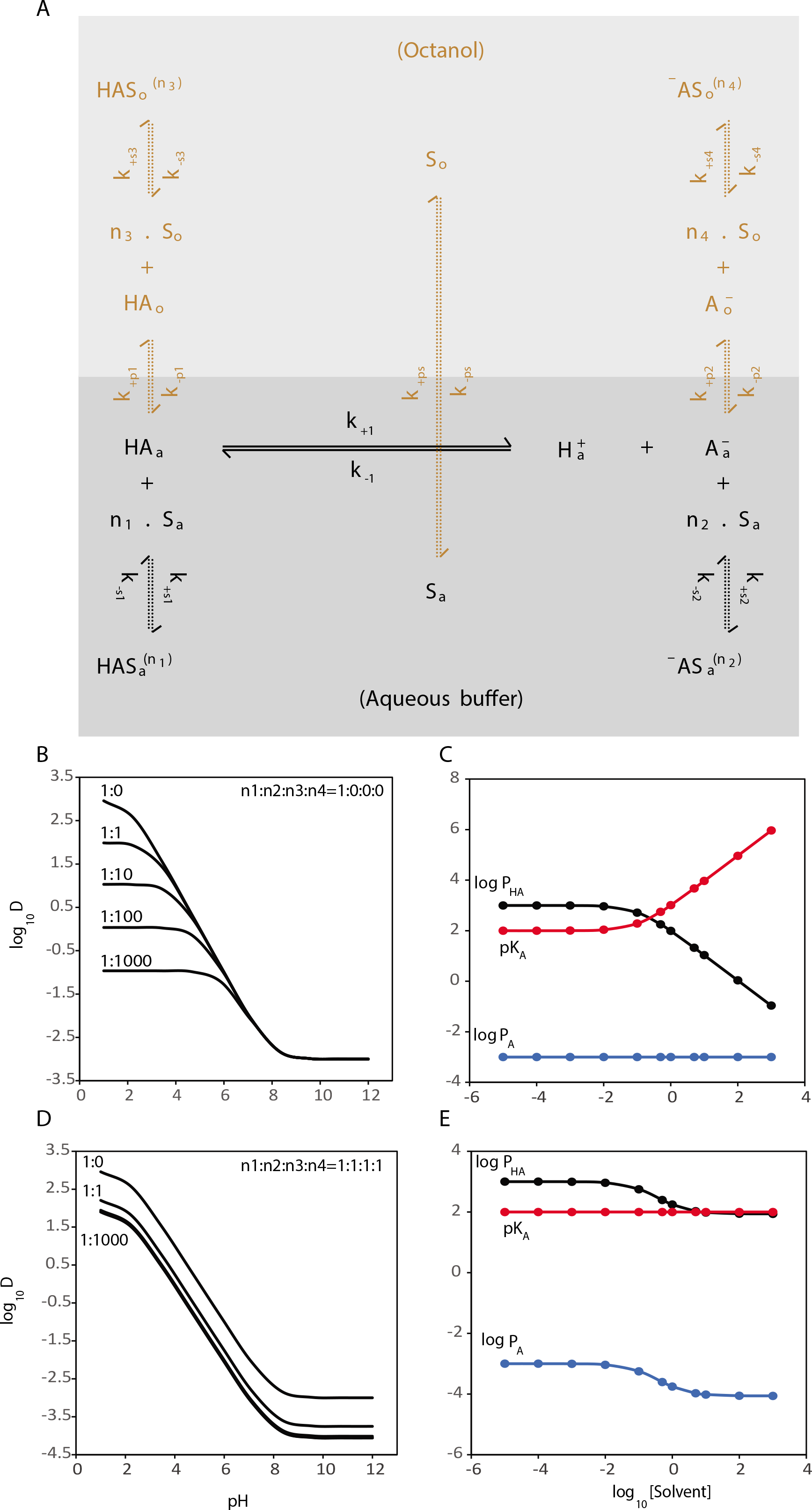
(A) The kinetic model for a monoprotic acid in the presence of a co-solvent ([S]). In this mechanism, there exists a reversible equilibrium between *HA_a_*, and 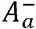 in the aqueous phase, whose species also get partitioned into octanol as *HA_o_*, and 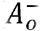 respectively. All four species, *HA_a_*, 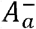, *HA_o_*, 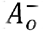 are co-solvated by the solvent, present in aqueous([*S_a_*]), or octanol phase ([*S_o_*]) with the stoichiometry of *n*_1_, *n*_2_, *n*_3_, *n*_4_, to form *HAS_a_*, ^−^AS_a_, *HAS_o_*, ^−^*AS_a_*, respectively. (B) & (D) Shows the *log*_10_*D* profiles of the monoprotic acid when the ratio of monoprotic to solvent is varied between 1:0 and 1:1000, with the stoichiometry of *n*_1_, *n*_2_, *n*_3_, *n*_4_, to be 1:0:0:0 for (B) and 1:1:1:1 for (D), respectively. (C) & (E) shows how the *log*_10_ *P_HA_*, *log*_10_ *P_A_*, *pK_A_* of the monoprotic acid varies as the solvent concentration increases, from 0 to 1000 (shown here in *log*_10_ scale). It is clear that the modulation of *log*_10_ *P_A_*, *log*_10_ *P_HA_* and *pK_A_* is strongly dependent on the degree of the co-solvation of different species. In the simulation, the *P_HA_*, *P_A_* and *K_A_* of the monoprotic acid were set to 10^3^, 10^−3^ and 10^3^ respectively, the concentration of monoprotic acid was set to 1 and the solvent concentration was set to 0, 10^−5^, 10^−4^, 10^−3^, 10^−2^, 10^−1^, 0.5, 1, 5, 10^1^, 10^2^, 10^3^ to simulate 12 different *log*_10_*D*, profiles. Only five of the solvent profiles corresponding to 0, 1, 10, 100, 1000 were plotted in (B) and (D). The simulated profiles were fitted to the monoprotic model, 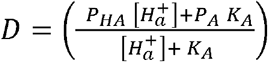 to obtain the apparent 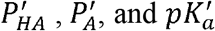 the logarithmic form of which are plotted against the logarithm of solvent concentrations.

In aqueous phase, the weak acid, [*HA*], exists in unionized form [*HA_a_*] and ionized form [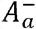]. The equilibrium between these two states can be written as (Eqn. 40),

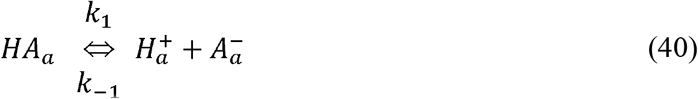

The distribution of a co-solvent between an aqueous buffer and octanol is given by (Eq. 41),

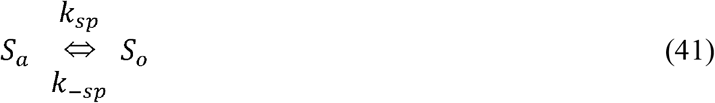

*k_sp_*, *k*_−*sp*_ are the forward and reverse kinetic rates for the partition of the co-solvent [*S*] between the aqueous (*S_a_*) and octanol phase (*S_o_*), respectively. The co-solvation of unionized form [*HA_a_*] and [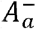] in aqueous buffer can be written as (Eq. 42 & 43)

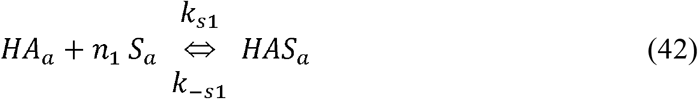

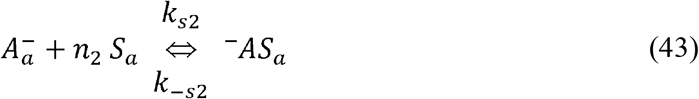

*k*_*s*1_, *k*_−*s*1_; *k*_*s*2_, *k*_−*s*2_; are the forward and reverse kinetic rates for the co-solvation of [*HA_a_*], [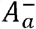] in the aqueous phase, respectively. *n*_1_, *n*_2_, are the number of solvent molecules required to co-solvate [*HA_a_*], [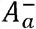], respectively.

[*HA_a_*], and [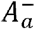], can be partitioned into octanol layer as shown below (Eqns. 44 & 45),

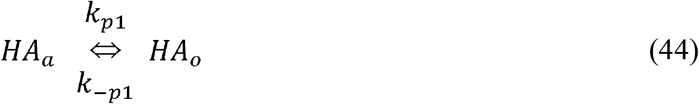

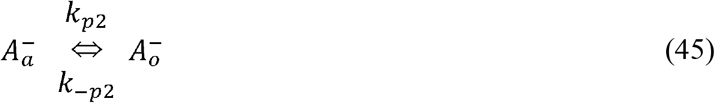

The co-solvation of unionized form [*HA_a_*] and [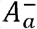] in octanol phase can be written as (Eqns. 46 & 47),

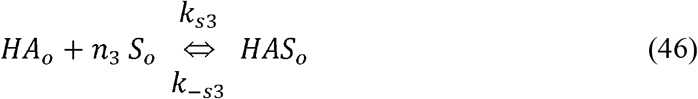

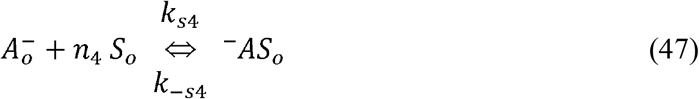

*k*_*s*3_, *k*_−*s*3_; *k*_*s*4_, *k*_−*s*4_, are the forward and reverse kinetic rates for the co-solvation of [*HA_o_*], [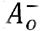] in the aqueous phase, respectively. *n_3_, n_4_*, are the number of solvent molecules required to co-solvate [*HA_o_*], [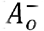], respectively.

#### 2.3.2. Algebraic method for monoprotic acid in the presence of co-solvent

Applying Le-Chaterlier’s principle, on the above equilibriums (Eqns. 40 - 47), and considering the law of conservation of mass for [*HA_T_*], and [*S_T_*], we solve for the analytical expressions of the ten species [*HA_a_*], [*HA_o_*], [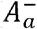], [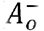], [*HAS_a_*], [^−^*AS_a_*], [*HAS_o_*], [^−^*AS_o_*], [*S_a_*] and [*S_o_*] (SI.6).

The distribution coefficient (D) is defined as (Eqn. 48),

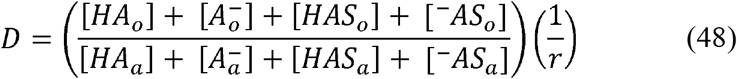

On substituting the analytical expressions for 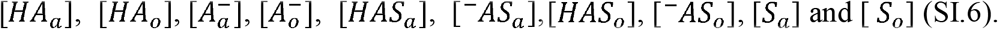 into Eqn. 48, we obtain Eqn. 49,

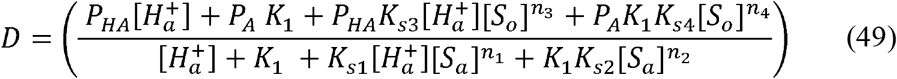

#### 2.3.3. Dynamic method for monoprotic acid in the presence of co-solvent

The differential equations for the rate of change of different species can be framed based on the kinetic mechanism given by (Eqns. 40 - 47). The rate equation for ten species, 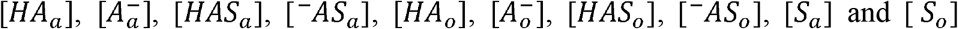, can be written as follows,

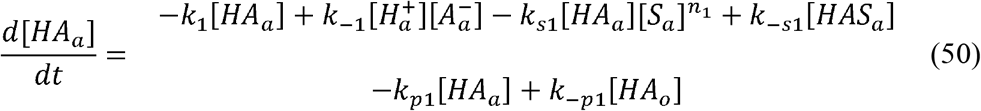

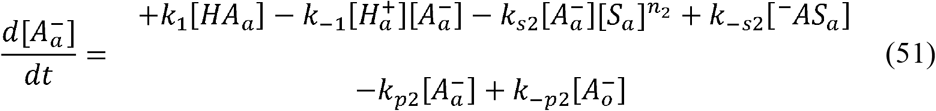

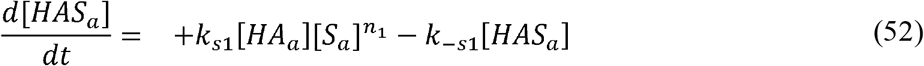

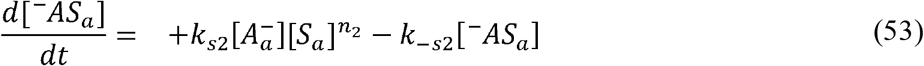

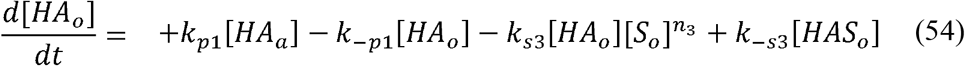

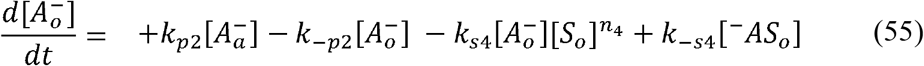

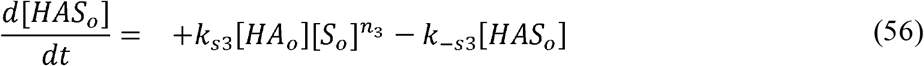

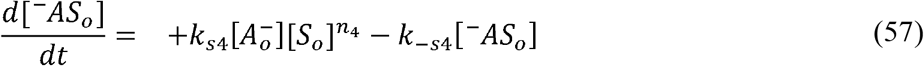

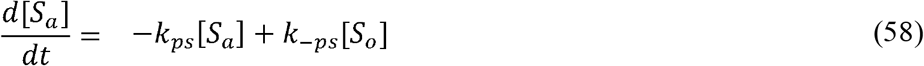

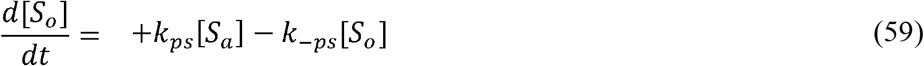

By numerically integrating the above set of coupled differential equations, Eqns 50 - 59, we obtain the concentration of ten species,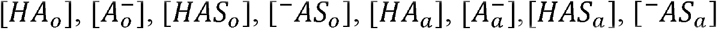 [*S*_a_] and [*S*_o_] at different time points. The resultant concentrations at equilibrium, (i.e. concentrations at *t* → ∞, or a long time period (>10^10^ *s*)), can be substituted into the Eqn. 48, to obtain the distribution coefficient. In the above model, [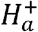] or *pH*, is assumed to be constant with respect to time. To simulate the *log*_10_*D* profile in Figure 3B&D, the parameters such as *k*_1_, *k*_−1_, *k*_*s*l_, *k*_−*s*l_, *k*_*s*2_, *k*_−*s*2_, *k*_−*s*3_, *k*_*s*3_, *k*_−*s*4_, *k*_−*s*4_, *k*_*p*1_, *k*_−_*p*_1_, *k*_*p*2_, *k*_−*p*2_, *k*_*pS*_, *k*_−*pS*_, were set to 10^−2^, 1.0, 10^7^, 1.0, 10^4^, 1.0, 10^8^, 1.0, 10^5^, 1.0, 10^1^, 1.0, 10^−3^, 1.0, 8* 10^−2^, 1.0 respectively. *n*_1_, *n*_2_, *n*_3_, *n*_4_, were set to 1,0,0,0, Figure 3B and *n*_1_, *n*_2_, *n*_3_, *n*_4_, were set to 1,1,1,1, Figure 3D. The initial concentrations of the eight species 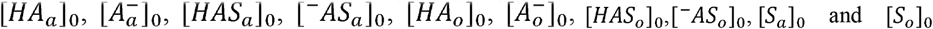 were initialized to 1.0, 0,0,0,0,0,0,0, [*S_T_*],0 respectively. A total of 12 simulations were performed with the following concentration of the total solvent, [*S_T_*] as 0, 10^−5^, 10^−4^, 10^−3^, 10^−2^, 10^−1^, 0.5, 1, 5, 10^1^, 10^2^, 10^3^ and the *pH*, was varied linearly between 1 and 12.

## 3. Results

In a recent *log*_10_*D* comparison study, several amphoteric molecules of pharmaceutical interest were analysed using both shake flask (octanol-buffer) and potentiometric method [10]. Among those molecules, the experimental data of nalidixic acid, mebendazole, benazepril, telmisartan were considered for our current analysis [10]. Though, nalidixic acid and mebendazole are amphoteric in nature, the *log*_10_*D* profile for both these molecules could be well represented through a monoprotic and monalkaline model, respectively (Figure 4A & B). Despite the fact that, benazepril is a complex amphoteric molecule with two amino group and one carboxylic acid group, its *log*_10_*D* data could be explained through a simple amino acid model (monoprotic-monalkaline) (Figure 4C). On the other-hand, telmisartan, required a complex monoprotic-dialkaline model to fit its experimental data (Figure 4D). The optimized *pK_a_*, *pK_b_* and *log*_10_*P* values were consistent with the previous studies and are summarised in Table 1 [10].

**Figure 4.**
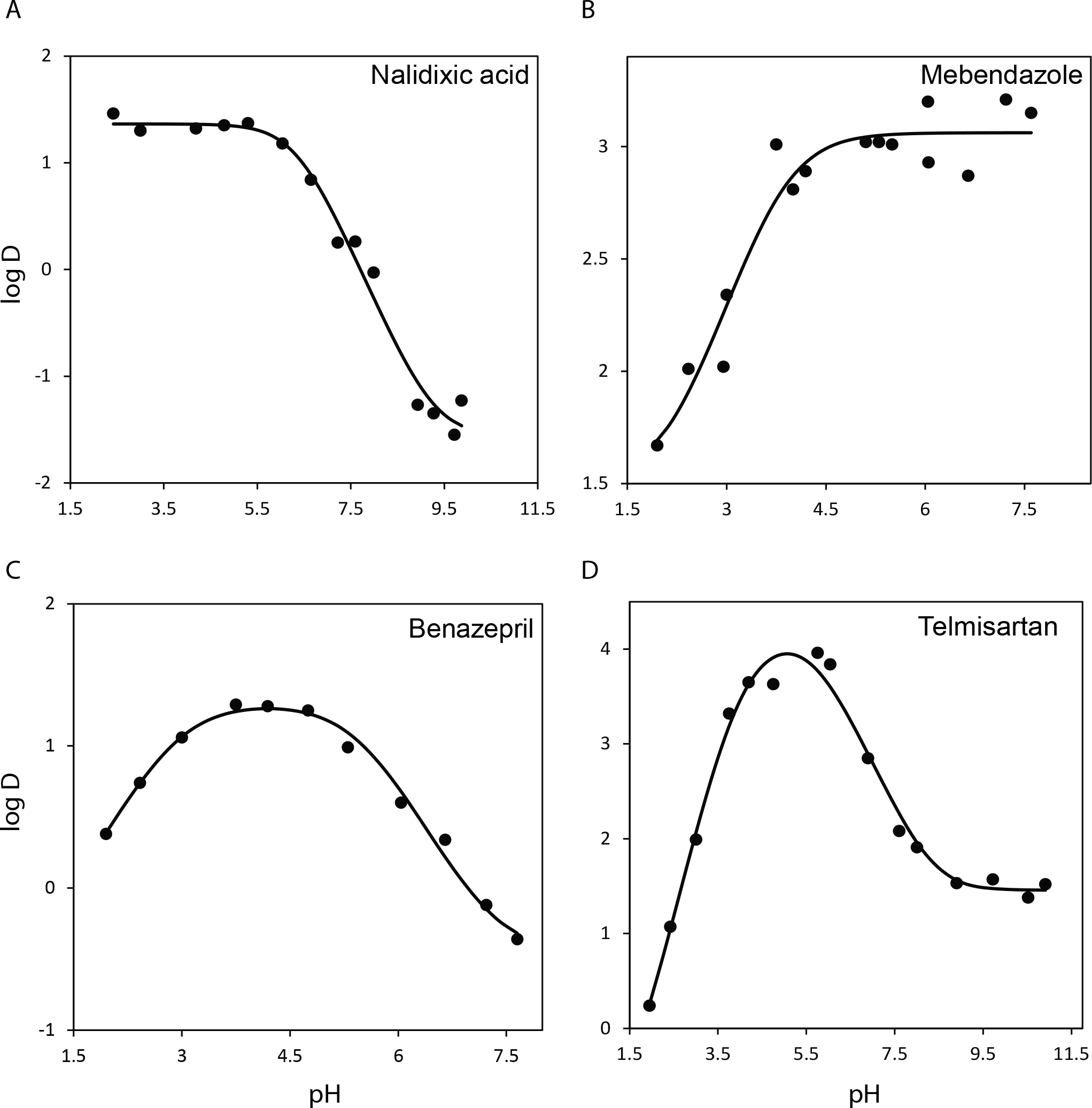
Experimental *log*_10_*D* analysis of (A) nalidixic acid, (B) mebendazole, (C) benazepril and (D) telmisartan using monoprotic, monoalkaline, simple amphoteric and diamino-monoprotic amphoteric models, respectively. The data were fitted using an inhouse written matlab code [16].

**Table 1.** Fit parameters for nalidixic acid, mebendazole, benazepril and telmisartan based on monoprotic, monoalkaline, simple amphoteric and diamino-monoprotic acid models, respectively. The parameters such as pK_A_, pK_B_, pK_B1_, pK_B2_ are equivalent to the conventional pK_a_which is related to the dissociation of the H^+^ion from the acid moieties. The pK_B_, pK_B1_, pK_B2_ are not the conventional pK_b_, which is related to the association of OH^−^ion with the base moieties.

|  | Nalidixic acid | Mebendazole | Benazepril | Telmisartan |
| --- | --- | --- | --- | --- |
| Model | monoprotic | monoalkaline | simple amphoteric | monoprotic-diamino amphoteric |
| Parameters |  |  |  |  |
| | $-1.363 \pm 0.0736$<br>($P_{HA}$) | $-3.059 \pm 0.0531$<br>($P_{BOH}$) | $0.458 \pm 0.1075$<br>($P_{HAB}, P_{-ABH^+}$) | $4.161 \pm 0.2749$<br>($P_{HABB}, P_{-ABBH^+}, P_{-ABH^+B}$) |
| Partition coefficients (P) | $1.714 \pm 0.1672$<br>($P_{A^-}$) | $-1.511 \pm 0.2575$<br>($P_{B^+}$) | $-1.307 \pm 0.0499$<br>($P_{HABH^+}, P_{-AB}$) | $1.458 \pm 0.0935$<br>($P_{HABBH^+}, P_{HABH^+B}, P_{-ABH^+BH^+}, P_{-ABB}$) |
| | | | | $-71825692.165 \pm 0$<br>($P_{HABH^+BH^+}$) |
| $pK_a/pK_b$ | $6.351 \pm 0.1261$<br>($pK_a$) | $10.248 \pm 0.1716$<br>($pK_b$) | $2.884 \pm 0.1046$<br>($pK_A$) | $5.6 \pm 0.405$ ($pK_A$) |
| | | | $5.481 \pm 0.1092$<br>($pK_B$) | $4.439 \pm 0.6262$ ($pK_{B1}$) |
| | | | | $3.608 \pm 0.3559$ ($pK_{B2}$) |

The simulation of the *log*_10_*D* profile of a monoprotic acid in the presence of salt was carried out to assess the effect of salt on *P_HA_*, *P_A_*, *pK_a_* values. The results show that, the addition of salt, affects only the *P_A_* value in a linear manner (directly proportional), whereas, the, *P_HA_* and *pK_a_* remained constant [5]. The effect of co-solvent (e.g. DMSO) on the *log*_10_*D* profile of a monoprotic acid was assessed through simulation. The *log*_10_*D* profile was primarily influenced by the differential ability of the co-solvent to co-solvate different species of monoprotic acid such as [*HA_a_*], [*HA_a_*], [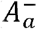] and [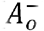]. For example, if the binding affinity (*K*_*S*1_, *K*_S2_, *K*_S3_, *K*_*S*4_) of the co-solvent ([*S*]) towards [*HA_a_*], [*HA_o_*], [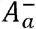], [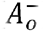], were assumed to be 1,1,1,1, respectively; and the stoichiometry *n*_1_, *n*_2_, *n*_3_, *n*_4_ to be either 1,0,0,0, or 2,0,0,0 or 3,0,0,0 respectively, then both *P_HA_* and *pK_a_*, were affected non-linearly, with increasing co-solvent ([*S_T_*]); whereas, *P_A_*, remained constant (Figure 4C). On the other-hand, if we let the stoichiometry *n*_1_, *n*_2_, *n*_3_, *n*_4_ alone to be varied as 1,1,1,1, respectively; *pK_a_*, remained constant; whereas, *P_HA_*, *P_A_*, varied non-linearly with increasing co-solvent concentration (Figure 4E) [3, 5]. Furthermore, if we let the stoichiometry *n*_1_, *n*_2_, *n*_3_, *n*_4_ to be 2,1,1,1, or 3,1,1,1, respectively, all the three parameters *pK_a_*, *P_HA_*, *P_A_*, varied non-linearly with increasing co-solvent concentration. Thus, we observe that the effect of co-solvent on *log*_10_*D* profile is non-linear, and is depended on the stoichiometry ratios that determine the degree of co-solvation of different monoprotic species.

## 4. Discussion

The complex *log*_10_*D* profile of telmisartan could be explained only through di-alkaline-monoprotic model. In this model eight different species were consider to exist in aqueous phase and an additional eight species were considered to be partitioned into the octanol phase. Considering only the eight species that are present in the aqueous phase, there are multiple ways a kinetic mechansim can be proposed by interconnecting these eight species among itself through a network of equilibrium reactions. For example, if we consider each species as a ‘node’ and the interconnecting equilibrium reactions as bidirectional ‘edges’, then the graph theory suggest a maximum of *N*(*N* − 1) = 56, edges or equilibrium reactions to exist among *N* = 8, species [14]. Under such a circumstance, developing an algebraic model based on the kinetic mechanism can be considerably simplified by choosing only seven unique equilibriums that inter connect any of these eight species. Further, by including the law of conservation of mass of the di-alkaline-monoprotic molecule as a constraint and an additional eight equations derived from the partitioning of the eight species between aqueous and octanol layer, we obtain 16 algebraic equations to solve for the concentrations of 16 species at equilibrium. On the other-hand, in the dynamic approach, the nature of the rate equation for the sixteen species will depend primarily on the network of equilibriums that is assumed to exist among the species. Even though the concentrations of the 16 species will vary during the pre-steady state condition, at equilibrium or steady state, the concentrations remains invariably the same, for equivalent kinetic mechanisms.

The model-fitting (Figure 4C, & D) was carried out using algebraic method (Eqn. 11 & SI Eqn. 155) and the resultant optimized parameters were used to simulate the *log_10_D* profiles through dynamic approach (SI Eqns. 156 - 171) The equilibrium concentrations obtained through algebraic and dynamic methods were comparable and equivalent to a degree of two decimal points for most of the data points (Table 2). The minor discrepancies seen in the accuracy of dynamic method compared to algebraic method is caused by errors in numerical integration when solving ordinary differential equations (ODE). If ODE’s are integrated over a long period of ‘integral time’, such as 10^8^s, the ‘calculation time’ required to solve the ODE’s and the ‘integration’ error, will increase significantly. This significant increase in time (> 2 hrs on a 8 cpu core personal computer) taken to solve the system of ODE’s is the major limitation in implementing dynamic approach within model-fitting routines to analyse the experimental data.

**Table 2.** Comparison of the log D values back calculated using algebraic and dynamic approach for Benazepril and Telmisartan, using simple amphoteric and diamino-monoprotic acid models, respectively. Accuracy to a degree of two decimal points was observed for most of the data points analysed through algebraic and dynamic approach.

| Benazepril |  |  |  | Telmisartan |  |  |  |
| --- | --- | --- | --- | --- | --- | --- | --- |
| *pH* | Experimental<br>($\log_{10}D$) | Calculated ($\log_{10}D$) | | *pH* | Experimental<br>($\log_{10}D$) | Calculated ($\log_{10}D$) | |
|  |  | Algebraic | Dynamic |  |  | Algebraic | Dynamic |
| 1.95 | 0.38 | 0.3876 | 0.3864 | 1.95 | 0.24 | 0.2541 | 0.2461 |
| 2.42 | 0.74 | 0.7380 | 0.7373 | 2.42 | 1.07 | 1.0431 | 1.0219 |
| 3.00 | 1.06 | 1.0643 | 1.0662 | 3.00 | 1.99 | 2.0612 | 2.0304 |
| 3.75 | 1.29 | 1.2420 | 1.2469 | 3.75 | 3.32 | 3.1856 | 3.1582 |
| 4.19 | 1.28 | 1.2624 | 1.2675 | 4.19 | 3.65 | 3.6251 | 3.6165 |
| 4.75 | 1.25 | 1.2278 | 1.2310 | 4.75 | 3.63 | 3.9105 | 3.9576 |
| 5.30 | 0.99 | 1.0945 | 1.0926 | 5.75 | 3.96 | 3.7741 | 3.7890 |
| 6.04 | 0.60 | 0.6794 | 0.6700 | 6.04 | 3.84 | 3.6019 | 3.6096 |
| 6.65 | 0.34 | 0.2187 | 0.2092 | 6.89 | 2.85 | 2.8998 | 2.8967 |
| 7.22 | -0.12 | -0.1469 | -0.1506 | 7.60 | 2.08 | 2.2678 | 2.2832 |
| 7.65 | -0.36 | -0.3177 | -0.3159 | 8.00 | 1.91 | 1.9612 | 1.9963 |
|  |  |  |  | 8.90 | 1.53 | 1.5632 | 1.5673 |
|  |  |  |  | 9.72 | 1.57 | 1.4747 | 1.4768 |
|  |  |  |  | 10.52 | 1.38 | 1.4597 | 1.4597 |
|  |  |  |  | 10.91 | 1.52 | 1.4580 | 1.4421 |

The simulation of the *log*_10_*D* profile of the monoprotic acid with salt, suggests an increase in apparent 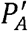, (i.e. *P_A_* in the presence of salt) with increase in salt concentration (*K_T_*) (Figure 3B & C). By comparing Eqn 29, with the simple monoprotic model (SI.1), we obtain 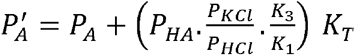, which shows that, 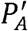 and *K_T_* are linearly related to each other, with the slope and intercept, 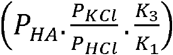 and *P_A_*, respectively (SI.9). The addition of salt, *KCl_T_*, tend to increase the formation of ion pair, *K*^+^: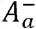, in the aqueous layer. The ion pair is formed between the ionized species 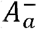 (from monoprotic acid) and the prevalently present cation *K^+^* (from *KCl*) [3, 5]. Unlike the charged species, 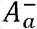, the neutral ion pair *K*^+^: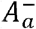 in aqueous phase can easily get partitioned into octanol as [*K*^+^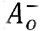], thereby, increasing the total concentration of [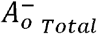] species (where, 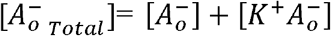 in the octanol phase). Since, 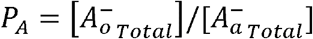 and is directly proportional to 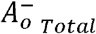, when 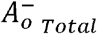 increases due to partitioning of the ion pair [*K*^+^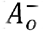], *P_A_* also increases proportionally.

To understand the complex *log*_10_*D* profile of the monoprotic acid in the presence of cosolvent we compare Eqn. 49, with the model of simple monoprotic acid and obtain the expressions for apparent 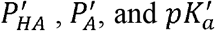 (SI.9) [3]. The apparent 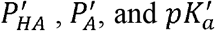 are the modified *P_HA_*, *P_A_* and *pK_a_*, of the monoprotic acid, respectively, in the presence of cosolvent. The resultant expressions, 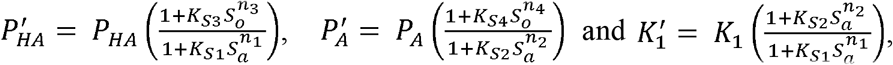, clearly shows that a non-linear relationship exist among 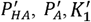 and the co-solvent concentration in octanol (*s_o_*) and aqueous (*s_a_*) phase. If we consider a simple stoichiometry of, *n*_1_, *n*_2_, *n*_3_, *n*_4_ to be 1 (or 2 or 3), 0,0,0; and the solvent binding affinities, *K*_*S*1_, *K*_*S*2_, *K*_*S*3_, *K*_*S*4_, to be 1,1,1,1, for the co-solvation of [*HA_a_*]:*S_a_*, [*HA_o_*]:*S_o_*, [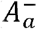]:*S_a_*, [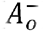]:*S_o_*, respectively, then, the apparent 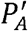 remains constant, i.e. 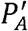 = *P_A_*, whereas, the apparent 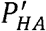 and 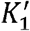 becomes non-linearly related to solvent concentration through the equations, 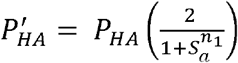 and 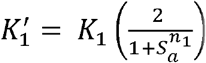, respectively (Figure 4C). For, *n*_1_ = 1, the apparent 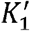, decreases non-linearly with increase in co-solvent concentration (i.e. 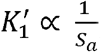). By definition, 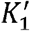 and *p*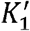 are inversely related to each other 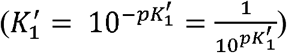. Therefore, we can infer that the apparent *p*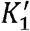 increases with increase in co-solvent concentration [15].

## Supporting Information

Supporting Information contains explicit derivation of log_10_D for mono-protic, di-protic acid, mono-alkaline, mono-protic acid with salt, mono-protic acid with solvent and monoprotic-dialkaline molecule. The expressions for apparent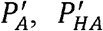 and 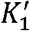 for monoprotic acid in the presence of salt and co-solvent are provided.

## Acknowledgments

JK would like to thank Jimma University (UN based faculty exchange program) and V ClinBio Pvt Ltd for their financial support; and Xavier Suburats (Universitat de Barcelona) for kindly providing the experimental data to carry out this work.

